# EndoAge Atlas: Non-linear dynamics of endothelial aging reveal organ-specific vascular trajectories in mice

**DOI:** 10.64898/2026.07.13.738166

**Authors:** Aisha Shigna Nadukkandy, Yonglun Luo, Lin Lin

## Abstract

Aging is a complex and multifactorial condition that leads to a gradual decline in organ functions and subsequent vulnerability to various diseases. Although the hallmarks of aging are well studied, the linear models developed based on these paradigms repeatedly fall short in capturing the dynamic, non-linear, and tissue-specific changes that occur because of aging. Furthermore, multi-organ level studies in humans are limited by the constraints of tissue availability and other confounding variables. This emphasises the importance of animal models in executing systemic investigations of aging. As the universal regulator of vascular function, endothelial cells (ECs) act as the initial responders to aging cues and undergo profound molecular and functional changes over time. This remodelling of the ECs is considered the central driver of multiple age-associated diseases. However, the global and tissue-specific molecular trajectories of aging in ECs are yet to be well characterised. Here, we introduce EndoAge atlas, a single-nucleus RNA-sequencing atlas comprising approximately 1.3 million ECs from 32 sex-balanced C57BL/6 mice across eight tissues and five distinct age groups. This atlas identifies tissue-specific endothelial heterogeneity, age-associated gene programs with distinct functional signatures, and uncovers non-linear gene expression trajectories across organs. Collectively, the EndoAge atlas constitutes a comprehensive reference for vascular aging across mammalian tissues, providing a foundational platform for developing targeted therapeutic strategies for age-related diseases.

## 1. Introduction

Aging is a physiological process characterized by the gradual decline of organ functions at cellular, tissue, and organismal levels, which increases the risk of developing nearly all diseases, including neurodegenerative diseases, cardiovascular diseases, and cancer. Over the years, studies focusing on aging have proposed several hypotheses on how the process manifests across different species [1–3]. Since the accumulation of DNA damage, stem cell exhaustion, and telomere shortening, along with other hallmarks of aging, are often implicated in the progression of aging, most studies have focused on identifying their linear relationship with age. However, aging is a multifactorial and dynamic process, and linear models often tend to oversimplify its molecular complexity. Consequently, non-linear trajectories of aging may better capture the intricate and time-dependent patterns of molecular dysregulation that occur across tissues. In this regard, several studies focusing on this non-linear trajectory at multi-omics levels have been published over the last decade [4–10]. Accordingly, these investigations have uncovered multiple molecular aging signatures that were previously unknown. However, the heterogeneous nature of aging across organs has made it challenging to fully elucidate its underlying mechanisms, especially in humans, partly due to the difficulty in acquiring multiple tissue samples from the same aging individuals for comprehensive cross-organ comparisons.

Hence, for cross-organ comparisons, animal models are often employed. Due to the ability to precisely control the genetic background, diet, sex, age, and environmental conditions in which these models are grown, animal models offer the opportunity to identify true biological aging effects, an approach that is often not feasible in humans due to several confounding variables, such as lifestyle and medications. Several efforts have been made to develop comprehensive reference atlases of mammalian aging. Recently, the PanSci atlas [11], a panoramic cellular characterization of aging in mice, with approximately 21 million cells from 13 distinct tissues, was developed using EasySci, an advanced single-cell combinatorial indexing strategy. Previously, the Tabula Muris consortium generated a murine single-cell atlas of 23 tissues, revealing 76 distinct cell types across six age groups [12]. A multi-omic atlas, encompassing data from epigenomics, transcriptomics, proteomics, and pharmacogenomics, has also been generated [13]. However, since aging studies demand substantial investment in time, resources, and animal maintenance, the generation of large-scale mammalian aging atlases still remains limited. In this regard, several short-lived model organisms, such as *Drosophila melanogaster* and *Caenorhabditis elegans*, which allow lifespan experiments to be completed within weeks, have become popular in aging studies, despite their limited genetic similarity to humans [14, 15]. However, studying all cell types across multiple organs poses significant technical and analytical challenges. Hence, the development of aging clocks, as well as studies, has primarily focused on ubiquitous cell types, such as immune cells. In fact, several aging clocks proposed over the years have been generated based on data from immune cells [16, 17].

One other ubiquitously present cell type that can be an ideal candidate in studying the systemic and organ-specific manifestation of aging would be the endothelial cells (ECs). Studies have shown that ECs undergo impairment of angiogenesis, extracellular matrix remodelling, inflammation, mitochondrial dysfunction, and oxidative stress, which ultimately remodel the vasculature during aging [18, 19]. As critical regulators of vascular function, the fact that endothelial senescence is related to most diseases makes it a suitable target for studying aging and its associated diseases. Moreover, the vasculature is also the primary sensor of systemic cues related to aging-associated factors, and thus the first-line responder to these cues. Also, some of the promising interventions in targeting vascular aging involve eradicating senescent ECs using Metformin, nicotinamide adenine dinucleotide (NAD+), hyperlipidemia-lowering agents, resveratrol, and senolytics [20]. Altogether, understanding endothelial response to aging is crucial for developing prevention strategies for aging-associated diseases, such as neurodegenerative and cardiovascular diseases.

Here, we present the EndoAge atlas, a single-nuclei RNA sequencing atlas comprising approximately 1.3 million ECs from 32 sex-balanced C57BL/6 mice, distributed across 8 major tissues, extracted from the acclaimed PanSci atlas. We were able to resolve the heterogeneous nature of vascular ECs in all these tissues and unravel several age-correlated genes across five age groups in all the tissues, and were able to characterise their functions. We were also able to identify a nonlinear pattern in gene expression changes across different tissues. Distinct molecules and functional pathways associated with these patterns were also identified. Altogether, our data is a comprehensive source for understanding endothelial aging in mammalian models across different tissues and could be used for further exploration of this matter.

## 2. Materials and Methods

### 2.1. Collection of publicly available data for snRNA sequencing analysis

For the development of the EndoAge atlas, we downloaded the h5ad file along with the processed count matrices, gene metadata, and cell metadata from the NCBI GEO website under accession number GSE247719. This data belongs to the PanSci atlas and is generated using the EasySci technique. EasySci is a cost-effective single-cell profiling technique developed as a modification of sci-RNA-seq3 [21, 22]. Unlike 10x Genomics, which relies on microfluidic droplets to isolate a barcode for each nucleus, EasySci employs a plate-based combinatorial indexing technique, which makes the workflow easier and highly scalable. This technique excels in throughput, making it ideal for generating massive cellular atlases, although it has lower per-cell transcript capture compared to 10x Genomics.

### 2.2. Data reproducibility checks and EC data extraction

The processed count matrix data obtained from the NCBI GEO repository were used to check the data quality and reproducibility of the PanSci data. The raw count matrix of each organ was reprocessed independently using a standardized pipeline in R (v4.3.1) using Seurat (v5.0.1) under Windows 11 (Europe/Copenhagen). For each organ, we generated Seurat objects from the sparse count matrix, calculated the percentage of mitochondrial genes, and plotted them alongside the number of UMI counts and gene counts to verify the distributional concordance with the PanSci atlas. Doublet cells were identified using the DoubletFinder (v2.0.4), and only singlets were retained for downstream checks. Gene biotypes were annotated by mapping the gene ENSEMBL IDs using the function useMart() to the ‘ensembl’ mart. This enabled stratified QC visualizations for protein-coding versus non-coding features, ensuring that the observed signal was not driven by non-coding artifacts alone. Furthermore, we extracted the singlet data and generated dimensional reduction plots using Uniform Manifold Approximation and Projection (UMAP). The reproducibility of the data was confirmed by the comparable cell yields from singlet cell selection and overlapping QC metric distribution across organs. Later, for convenience, we extracted the h5ad data, generated Seurat objects for each tissue, and subsetted ECs using the available metadata information and classical markers.

Subsequently, potential batch-specific variations were investigated within each organ prior to downstream analysis. The generated Seurat objects were normalized using the LogNormalize method, where the feature count from each cell is divided by the total counts for that cell and then multiplied by the scale factor, which is further log-transformed using log1p. Normalized data is used to identify the top 7,000 variable genes using the ‘vst’ method, and then scaled before applying principal component analysis (PCA) (by identifying the top 50 PCs), UMAP, and t-distributed Stochastic Neighbor Embedding (t-SNE) for dimensional reduction. The decision to select 7,000 variable genes is heavily influenced by the sequencing depth of the EasySci method (9,000 compared to 20,000 for the 10x Genomics method). Furthermore, the total number of ECs per organ was quantified and visualized using bar plots, while UMI distributions were compared across organs using boxplots, highlighting differences in sequencing depth and transcript capture efficiency (**Supplementary Fig. 1**).

### 2.3. Endothelial subclustering and heterogeneity detection

To identify and annotate EC subtypes within each organ, we first mapped the ENSEMBL IDs to MGI symbols using the biomaRt (v2.62.1) package. The Seurat object was further subclustered and annotated. Marker genes for each cluster were calculated (FindAllMarkers, Wilcoxon rank sum test using the “RNA” assay) and screened for coherent enrichment of canonical marker genes, to annotate artery (*Fbln5, Sema3g, Tm4sf1, Alpl*), capillary (*Ablim3, Kdr, Ccdc85a, Nrp1*), angiogenic (*Apln, Esm1, Kit, Nid2*), vein (*Vwf, Selp, Nr2f2, Plvap*) and lymphatic (*Prox1, Mmrn1*) EC subtypes. Cell types expressing both capillary and venous markers were annotated as capillary venous. Other specialised EC subtypes specific to each organ were annotated based on the markers available from the literature. At this point, cell types that expressed non-EC marker genes were removed. We also removed ECs belonging to lymphatic EC clusters, to avoid the confounding effect due to their distinct transcriptional regulatory program. From the 13 tissues extracted and processed from the PanSci atlas, the data from brain, colon, ileum, jejunum, and duodenum were removed from further analysis based on quality and relevance assessment. For brain data, the dimensionality reduction plot obtained for the ECs was inconsistent with known vascular endothelial signatures available in literature, indicating annotation artifacts. The data from colon, ileum, jejunum, and duodenum exhibited strong enrichment of lymphatic endothelial cells, with minimal or absent expression of vascular ECs. Thus, this data was omitted from further analysis to ensure a comparative aging framework across organs. Post data processing, we obtained a total of 1,295,123 cells.

### 2.4. Detecting linearly changing molecules using linear regression analysis

To quantify the gene-associated changes associated with aging, we performed organ-wise linear regression analysis. Within each organ, single-nuclei data were aggregated at the biological replicate level using the AggregateExpression() function to obtain pseudobulk expression profiles per mouse ID. This decision was made because the EasySci method is developed using low sequencing depth, resulting in a sparse gene-cell matrix data with excessive dropouts. Consequently, the subsequent cell-level analysis is statistically unstable and underpowered. Thus, all the analysis presented in this manuscript is based on pseudo bulk data (per mouse ID, unless otherwise specified). Age was encoded as a continuous numerical variable, and a linear model was fit using ordinary least squares to estimate the age-associated slope, the standard error, as well as the corresponding p-values. To avoid the potential inflation due to low sample size, we implemented a permutation-based test (n = 1000) to randomly shuffle the age values. The proportion of Beta.Estimate values exceeding the observed changes were used to identify the empirical p-values and false discovery rate across all genes for each organ. The data was exported as Excel files for cross-organ comparisons.

### 2.5. Detecting linearly changing molecules using Spearman correlation analysis

To confirm our findings using linear regression analysis, we employed Spearman correlation to endothelial pseudo bulk data from each of the 8 tissues. For every gene, the Spearman rank correlation coefficient (rho.rho) was calculated between the gene expression and age. To evaluate the statistical significance, we employed the permutation-based approach (n=1000), where age values were randomly shuffled. Empirical p-values were calculated as a fraction of permuted rho.rho values exceeding the observed effect, a multiple test correction was done using the Benjamini-Hochberg method. Genes with FDR<0.05 were considered to be significantly correlated with age. To visualise the genes that are significantly enriched across all organs, UpsetR (v1.4.0) package was used. The results obtained for finding linearly changing molecules using both methods were compared by running a Spearman correlation test between the results from the linear regression and Spearman correlation, after rescaling the Beta.estimate value to exist within the scale of 1 and -1.

### 2.6. Detecting non-linear changes using differential gene expression analysis

To trace the transcriptional remodelling across ages within each organ, we performed organ-wise differential gene expression analysis (DEG) between consecutive ages (ie; 3 to 6, 6 to 12, 12 to 16, and 16 to 23 months) using the top 7000 variable genes. DEGs were computed using the function FindMarkers() which uses Wilcoxon rank-sum test, and genes were considered to be significantly deregulated if they have an absolute log2FC >/= 0.25 and adjusted p-value < 0.05. To visualise this dynamic transition, alluvial plots were generated using the ggalluvial (v0.12.5) package.

### 2.7. Differential Expression Sliding Window Analysis (DE-SWAN)

To identify the non-linear gene expression wave across different ages in each organ we employed the Differential Expression Sliding Window Analysis (DE-SWAN) on the pseudobulked endothelial transcriptome. Within each organ, snRNA seq data was aggregated at the biological replicate level using the AggregateExpression() function to obtain pseudobulk expression profiles per mouse ID. For each organ, sliding windows of one age group on either side were constructed and defined as ‘left’ and ‘right’ sample sets when compared to the centre age. Within each window, differential expression was tested between the age groups on either side using a limma-voom model, where voom normalization with TMM factors and empirical Bayes moderation is done. Significant genes were identified based on the criteria that it has an absolute log2FC >/=0.25 and FDR < 0.05. Each gene was then classified based on directionality ie; “Up_in_Right”, “Up_in_Left”, or “Non_sig”, and the number of significant genes per window was summarized to quantify local amplitude. Furthermore, for statistical significance, we employed the permutation-based approach (n=1000), where age values were randomly shuffled for each mouse and empirical p-values were calculated. Windows exhibiting a local maximum (crest) in the number of significant genes were considered as an inflection point in EC aging for that organ.

### 2.8. Gene Ontology (GO) enrichment analysis

GO enrichment analysis was performed using the package clusterProfiler (v4.14.6). Biological pathways with average log2FC > 0.25 and FDR < 0.05 were considered to be significantly enriched.

### 2.9. Data visualisation

All visualizations were generated using R (v4.3.1) using Seurat (v5.0.1) and ggplot2 (v4.0.0) and were curated using Inkscape and Biorender.

## 3. Results

### 3.1. EndoAge Atlas: Multi-organ map of distinct EC populations during aging

To gain a comprehensive understanding of ECs during aging, we extracted single-nuclei RNA sequencing data from the PanSci atlas, which was generated using an advanced single-cell combinatorial indexing strategy termed EasySci [11, 21]. Data extraction was followed by strict quality control processes and selection of ECs based on the canonical marker genes. The global object consisted of approximately 1.3 million ECs across eight tissues (brown adipose tissue (BAT), stomach, kidney, perigonadal adipose tissue (gWAT), muscle, inguinal adipose tissue (iWAT), liver and heart) from 32 sex-balanced C57BL/6 wildtype mice across five age groups of 3, 6, 12, 16, and 23 months that represent the human age of 20, 35, 42, 50 and 68 years, respectively (**Fig. 1A-C**). Heart tissue contained the highest number of ECs (587733), whereas the stomach had the least (33627) (**Supplementary Fig. 1D**). An average of 906 different transcripts was detected per cell (maximum, 1140 in the heart and minimum, 616 in the liver) (**Supplementary Fig. 1C**).

**Fig. 1:**
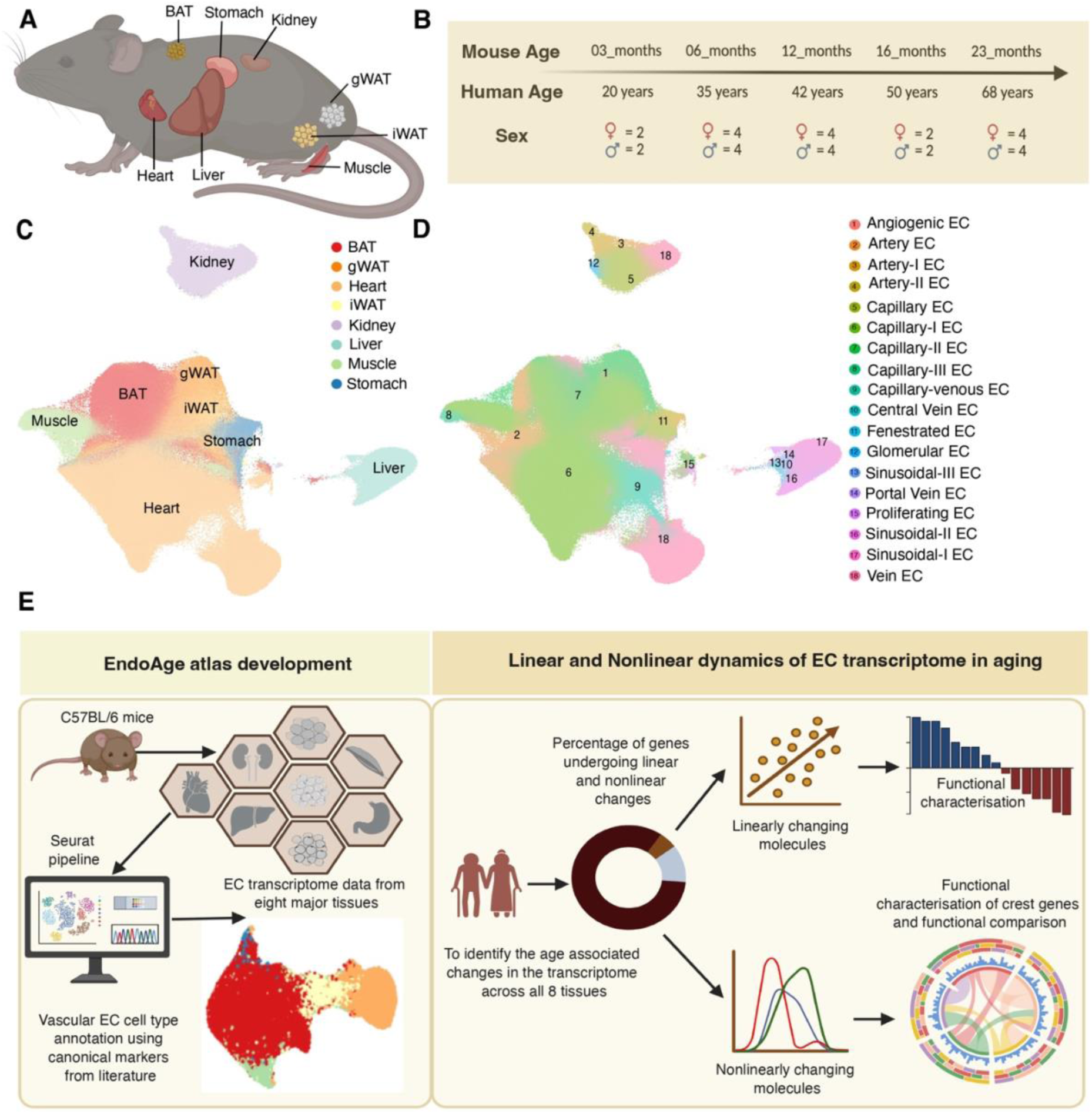
Overview of the experimental design and major cell types annotated in the EndoAge atlas. A) Schematic representation of the m,ajor tissues used to develop EndoAge atlas. B) Schematic representation of the age and sex distribution of the mice used for the experiment. C) An aggregated UMAP representation of all the EC data extracted across all tissues. D) An aggregated UMAp representation of the major cell types annotated across all tissues. E) Schematic illustration of the manuscript outline of the EndoAge atlas.

To characterize cellular heterogeneity across the tissue, we annotated the ECs according to the major cell types present throughout the vascular tree. We observed a consistent presence of capillary, vein, and artery ECs in all the tissues (**Fig. 2A-B**). Cell types that expressed canonical markers of both capillary and vein were termed capillary-venous and are likely to represent venules [23]. As reported previously, capillary ECs were more abundant in all tissues than artery, vein, and lymphatic ECs [23]. Capillary ECs that expressed *Top2a*, *Hmgb2*, and *Stmn1* were annotated as proliferating ECs, and those that expressed known tip-cell markers like *Apln*, *Nid2*, and *Kit* were termed angiogenic ECs. *Plat, Tbx3*, and *Tspan7* expressing capillary ECs in the kidney were annotated as glomerular ECs. As reported previously, we identified fenestrated ECs expressing *Col13a1, Plvap*, and *Igfbp7* in iWAT, but not in BAT and gWAT [24]. We were also able to annotate liver ECs into sinusoidal, central vein, and portal vein ECs [25]. All together we identified three different subtypes of sinusoidal ECs. Lymphatic ECs expressing *Lyve1, Mmrn1*, and *Prox1* were also identified but were excluded from further analysis to avoid the confounding effect due to their distinct transcriptional regulatory program. In line with the previously reported EC atlas in mice and humans, we observed transcriptional diversity in EC populations across tissues [23, 26].

**Fig. 2:**
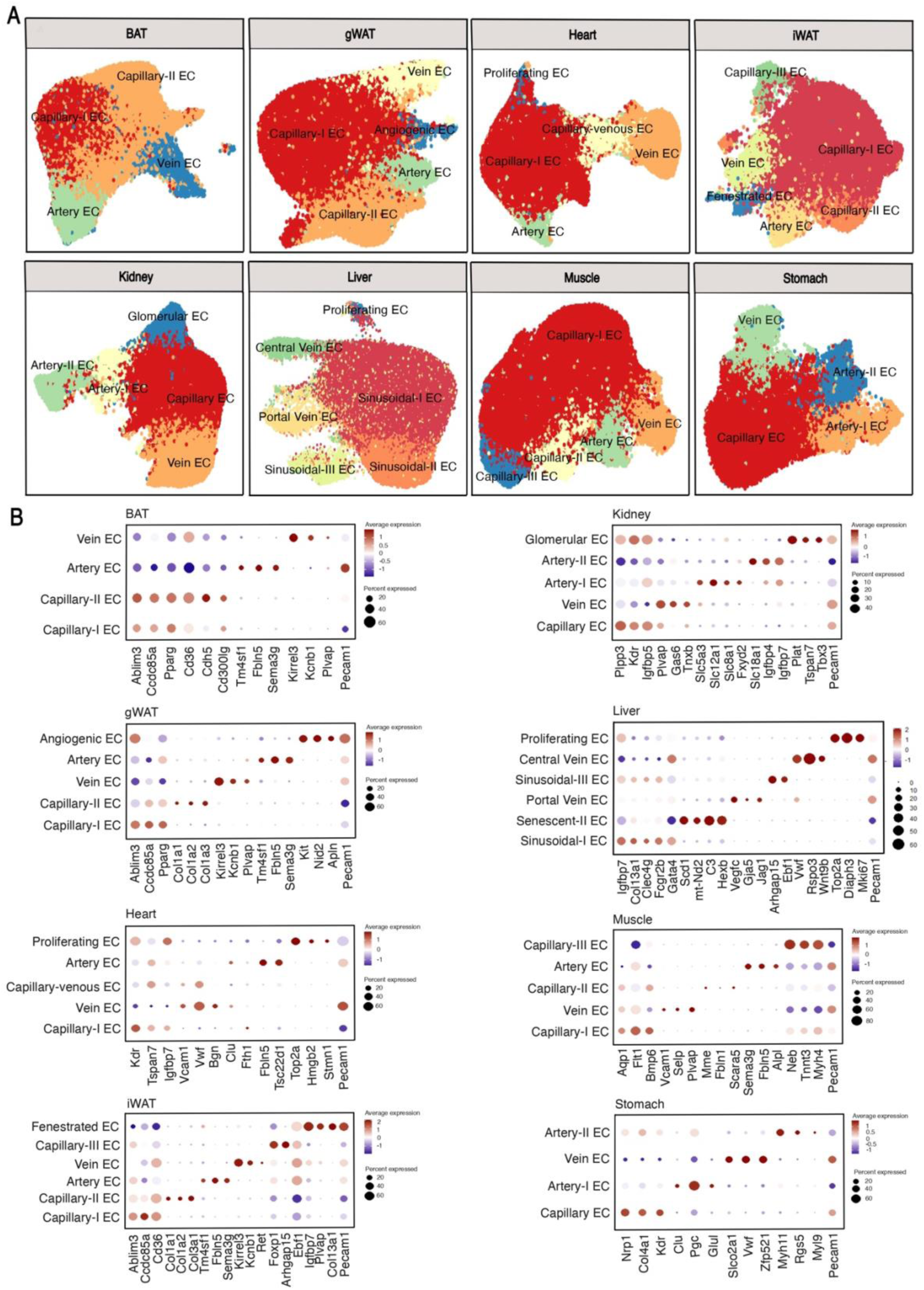
Cross-tissue annotation of EC subtypes and the markers used for annotation. A) UMAP visualization of major EC subtypes annotated across all eight tissues. Broad endothelial populations, including capillary, venous, and arterial ECs, are conserved across tissues. B) Dot plot showing the expression of canonical marker genes derived from the literature and used to annotate the major EC subtypes across all tissues. Dot size represents the percentage of cells expressing each gene, and color intensity indicates the scaled average expression level.

Furthermore, a gradual phenotypic zonation of ECs was observed across all tissues, with ECs arranged along the arteriovenous axis, with arterial ECs positioned at one end, venous ECs at the other, and capillaries and venules occupying intermediate states (**Fig. 2A**). This vascular tree hierarchy was also observed after data merging in all tissues, indicating that the transcriptional basis of vascular heterogeneity is conserved across all tissues (**Fig. 1C-D**).

### 3.2. Diverse functional roles of age-associated genes across organs

To identify the molecules that change linearly with age, both linear regression and Spearman correlation, used to detect linearly changing molecules, were applied separately to the transcriptomic data from each tissue. Both approaches yielded highly consistent results; however, Spearman correlation was used for further analysis (**Fig. 3A**). Notably, only a small fraction of genes exhibited linear correlation with age across all tissues. The greatest proportion was observed in gWAT, where 30.51% of genes showed age-associated linear changes, followed by the liver with 10.8%. In contrast, all other tissues displayed less than 10% of genes correlating linearly with age, with the stomach showing the lowest proportion at 1.81% (**Fig. 3B-C**). To corroborate our findings, we employed the permutation approach, which yielded consistent results. Subsequently, these age-associated genes were subjected to cross-tissue comparison, revealing that no genes consistently correlating with age are commonly present across all tissues. This might be an indicator of tissue specificity in age-related transcriptional programming (**Supplementary Fig. 2**)

**Fig. 3:**
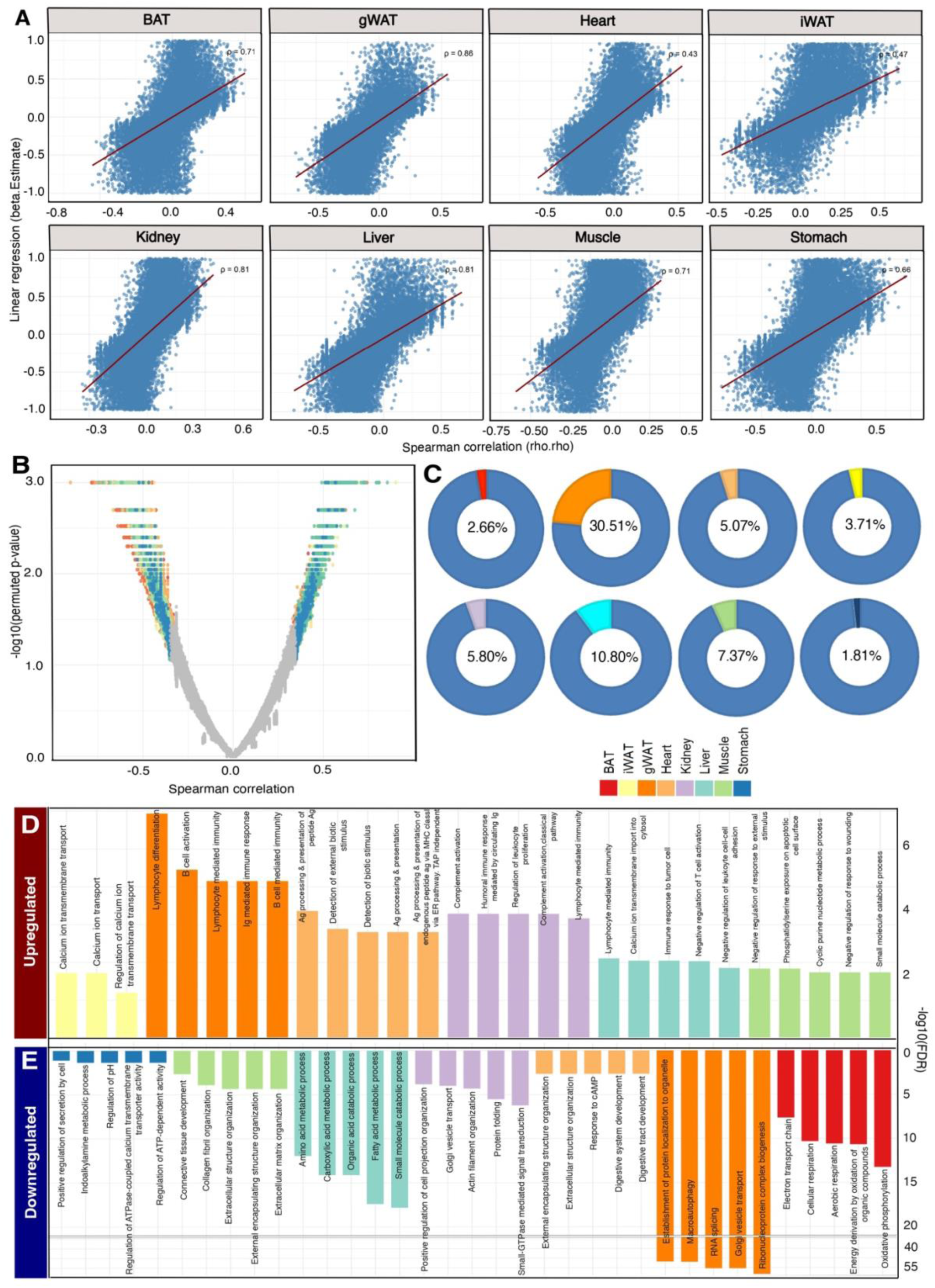
Identification of linearly changing molecules during aging. A) The Spearman correlation is highly correlated to the beta coefficients from linear regression models for the transcriptomic data from all tissues. B) Volcano plot showing significantly changing molecules with age identified using Spearman’s correlation (pval < 0.05). C) Pie chart showing the percentage of genes that are linearly correlated with age in each tissue. Barplot depicting selected pathways after enrichment of genes that are D) positively and E) negatively correlated with age.

The functional characterization of these linearly changing molecules further reinforced this observation. Gene ontology enrichment of these linearly correlated genes revealed high functional heterogeneity in how these genes are translated across different organs. While comparing genes negatively correlated with aging across different organs, BAT tissue exhibited downregulation in genes involved in cellular respiration-related GO terms, including aerobic respiration, oxidative phosphorylation, and the electron transport chain. Studies have reported a reduction in BAT mass and activity during aging in both rodents and humans, which is often attributed to mitochondrial dysfunction in aging BAT tissues, similar to what we observe here [27, 28]. Cellular protein synthesis capacity was compromised in gWAT, indicated by a decrease in genes responsible for RNA splicing, Golgi vesicle transport, and ribonucleoprotein complex biogenesis. The response to cAMP and the ability of heart ECs to organize the extracellular matrix are observed to decrease with aging. Kidney-deregulated genes were related to small GTPase-mediated signal transduction. The gradual decrease in small GTPase signalling was also accompanied by actin filament reorganization, indicating a disruption to the physiology of kidney ECs during aging, since this signalling pathway is known to control cell proliferation, differentiation, apoptosis, and cellular growth in the kidney [29]. Interestingly, liver ECs underwent severe metabolic reprogramming when compared to other organs. Several metabolic processes, including amino acid, carboxylic acid, and fatty acid metabolism, were observed to be disrupted gradually during aging, along with organic acid and small-molecule catabolic processes. ECs in muscle exhibit declining collagen fibril and extracellular matrix organizational function with age, whereas stomach ECs appear to contribute to the decline in secretory function within the organ. A decrease in indoalkylamine metabolism is accompanied by a decrease in pH regulation and regulation of ATPase-coupled calcium transmembrane transport in stomach ECs with age (**Fig. 3E**).

However, comparing genes positively correlated with aging across different organs (**Fig. 3D**), we observed that ECs in iWAT exhibited increased calcium ion transport ability, a characteristic usually observed in ECs during vascular contractility [30]. Muscle ECs enriched pathways that are responsible for stimulus-response. However, ECs in most tissues, such as gWAT, heart, kidney, and liver, were observed to exhibit a gradual increase in their immunomodulatory roles. GO terms related to lymphocyte-mediated immunity and differentiation, especially for B cells, have been increased in gWAT, whereas several antigen presentation and processing-related GO terms were increased in Heart ECs. Complement activation function is increased in kidney ECs, which has also been reported previously and could be correlated to age-associated kidney decline and associated chronic inflammation [31]. In the liver, a decrease in metabolic capability was followed by an increase in lymphocyte-mediated immunity. This trend aligns with the concept of inflammaging, suggesting that ECs progressively adopt pro-inflammatory and immune-regulatory phenotypes with age, thereby contributing to the chronic inflammatory characteristic of aging tissues.

### 3.3. Non-linear dynamics of aging across different organs

Next, we employed the Wilcoxon test to compare each age group with the consecutive ones, which revealed that a substantial proportion of genes have undergone non-linear changes across all organs. The Sankey plot, illustrating genes that are significantly deregulated (after adjusting p-values using the Bonferroni correction method, adjusted p-value < 0.05) in at least one age comparison, highlights that endothelial aging follows a non-linear trajectory (**Fig. 4A**). Recent longitudinal studies in humans have also stated that this non-linear nature exists at the multi-omics level during aging [10]. Non-linear gero-markers of the aging cardiac system have also been identified recently in mouse models [32]. To our knowledge, this is the first study to identify non-linear dynamics of aging in ECs in multiple organs in mice.

**Fig. 4:**
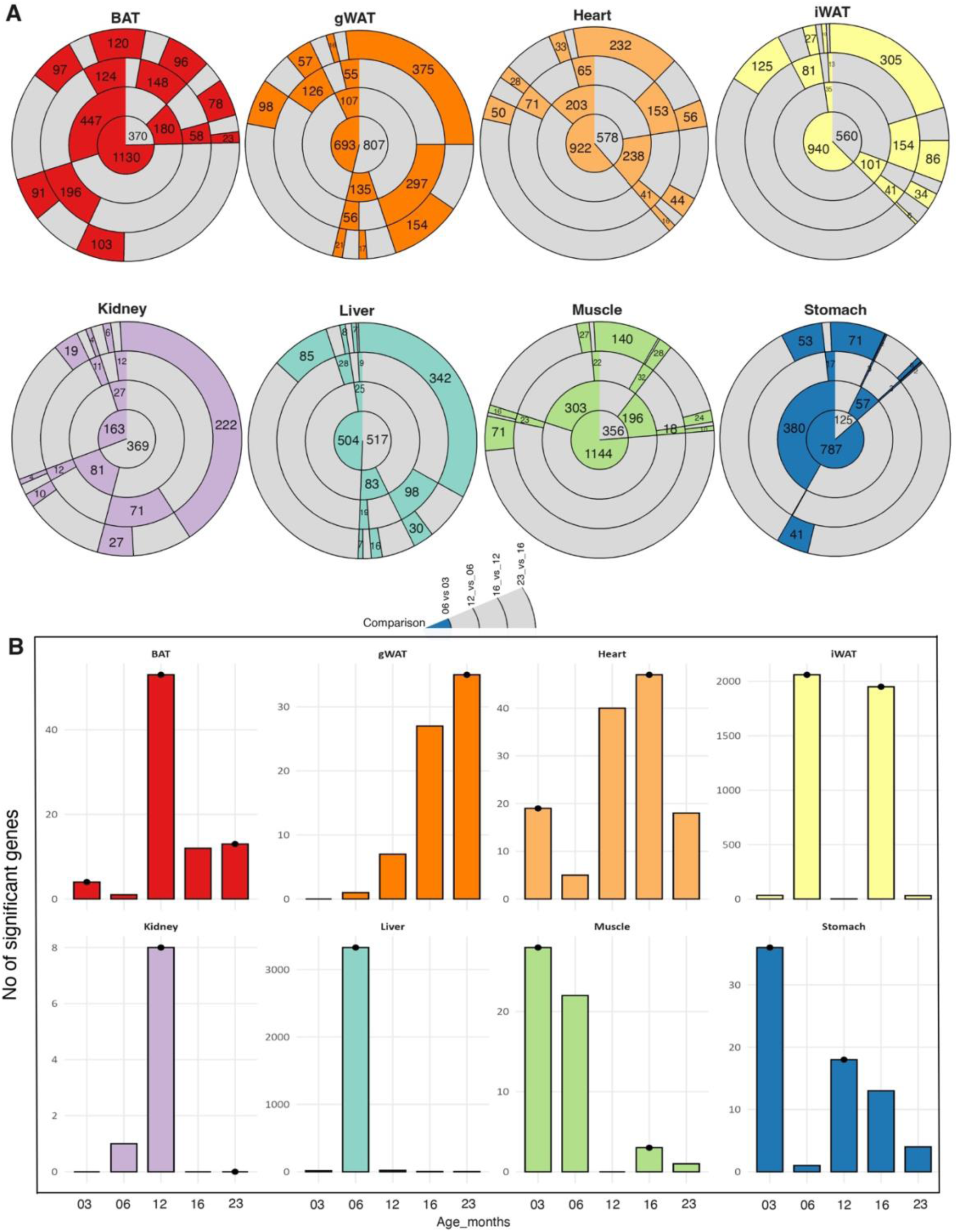
Identification of the non-linear dynamics of aging: A) Sankey plots illustrating the number of differentially expressed genes (Wilcoxon’s test, adjusted p value < 0.05) identified by comparing each age group with its consecutive age group across all tissues, revealing nonlinear patterns of gene expression changes over time. Coloured ribbon determines the significant genes, and grey ribbons indicate the non-significant genes. B) Bar plots showing the number of genes significantly associated with age in each tissue, as identified using the DESWAN method. Bars marked with a dot indicate ages at which a crest was detected, representing points of pronounced transcriptional change.

To screen the wave of gene deregulation during aging, we employed differential expression sliding window analysis (DE-SWAN) on the top 7,000 variable genes per organ. This algorithm identifies deregulated molecules within a window, performing pairwise comparisons between adjacent age groups as the window slides sequentially across the ages from young to old. This revealed substantial changes that occur at different ages in multiple organs, reiterating the non-linear aging theory and highlighting heterogeneity in how this non-linearity is manifested across different organs. A single crest was observed in gWAT, kidney, and liver, but at different age stages. Confirming the results of the Spearman correlation test, the DESWAN analysis revealed a consequent increase in the number of significant genes in gWAT (**Fig. 4B**). The functional characterisation of the crest observed at the 23 months shows that these genes are involved in responding to various stimuli (**Fig. 5C**). Notably, we observed changes in the expression of genes such as *Col3a1, Col5a1, Col1a1, Col1a2,* and *Col6a1*, indicating the reorganization of the extracellular matrix (ECM) and alterations in collagen fibril organization (**Supplementary Fig. 4**). This biological shift is expected, as ECs in adipose tissue are known to become fibrotic, characterized by increased ECM deposition and tissue stiffening [33]. Kidney ECs exhibit a distinct shift over 12 months, which is predominantly characterized by changes in genes related to immune system processes. Several pathways involved in the complement cascade, B-cell-mediated immunity, and humoral immune responses show significant alteration in mice at this age. Like gWAT, the genes that underwent abrupt changes in the liver were related to stimulus response. The liver exhibited a shift in the expression of over 3,000 genes at 6 months of age, and surprisingly, many of these genes belonged to the olfactory receptor (OR) and G-protein-coupled receptor (GPCR) categories. OR genes are being increasingly studied for their role in liver metabolism [34–36]. We also observe changes in immune-related genes, such as *Tlr6, Tlr9,* and *Trem2*, at this stage in the liver (**Supplementary Fig. 5**).

**Fig. 5:**
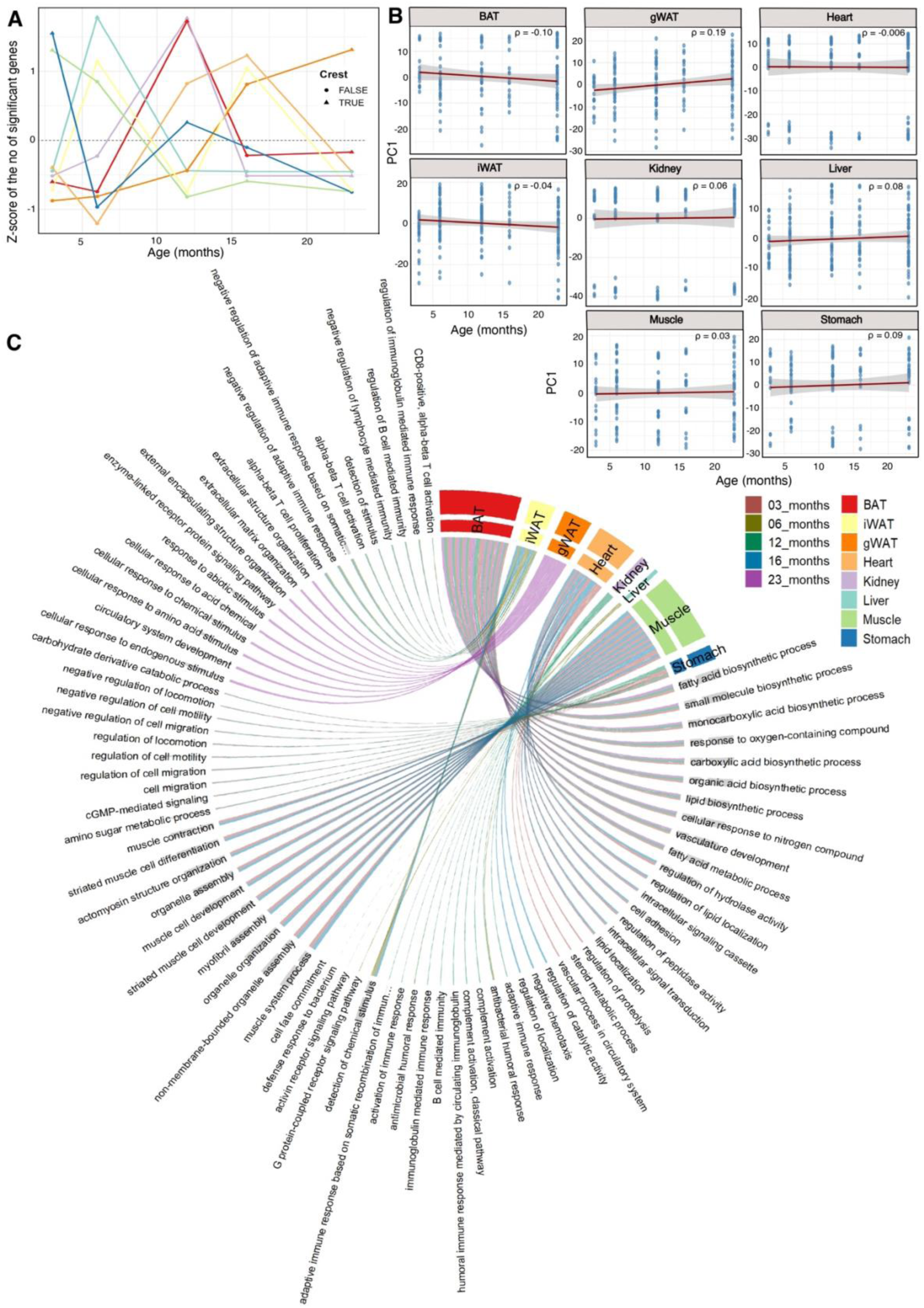
Functional characterisation of the nonlinear genes during aging: A) Line plot depicting the DESWAN results (Z-score normalised), indicating that ECs in each organ respond to aging in a different way. B) Scatterplot depicting the Spearman correlation between the first principal component and ages for the transcriptomic data from ECs of each tissue. The shaded area around the regression line represents a 95% confidence interval. C) A comprehensive circos plot showing the functional characterisation of the genes belonging to each crest within the respective tissue.

Two crests were identified in the muscle, stomach, iWAT, and heart ECs. Among these muscles and stomach ECs displayed a gradual decrease in the number of significant genes with age. Genes belonging to both the crests in muscle ECs were involved in muscle cell development, actinomyosin structure organisation, and cellular morphogenesis. Interestingly, these genes were upregulated at 3 months, while they were completely downregulated at 16 months, indicating muscle growth at a young age, and a rapid decline in this process as the mice reached 16 months of age. In stomach ECs, both the crests involved genes related to cellular migration and motility. ECs belonging to the iWAT tissue, which form crests at 6 and 16 months, are involved in genes associated with T-cell immunity and stimulus response. The gene expression levels increased significantly within 6 months and then exhibited an abrupt decrease by 16 months, indicating the possibility of adaptive immune-vascular interaction at the adult age of 6 months, along with the possibility of immunosuppression and vascular senescence at age 16 months. Heart ECs follow the wave through two crests, one at 3 months and the other at 16 months. At the earlier stage, genes are increased in expression for lipid localization and protein-lipid complex remodelling, while in the latter, genes are mostly downregulated and are involved in hydrolase activity, lipid localization, and chemotaxis. BAT is the only tissue with three crests with genes involved in fatty acid biosynthesis, small molecule biosynthesis, and monocarboxylic acid biosynthesis. Altogether, our results depict nonlinearity in how gene expression changes occur during aging in ECs across all analysed tissues. This is also confirmed by plotting the first principal component, which depicts the direction of maximum variability in the pseudo bulk (per mouse and cell type) gene expression data, which, when plotted along the age groups, exhibited no correlation (**Fig. 5B**). Notably, we observed substantial variability in PC1 values across cell types, indicating that cell-type– specific differences contributed more to the observed variance than aging (**Supplementary Fig. 3**).

## 4. Discussion

We present the EndoAge atlas, a comprehensive endothelial transcriptome atlas comprising approximately 1.3 million cells across eight different tissues in mice. Using this time series panel transcriptome dataset, we unraveled the heterogeniety of vascular ECs in all these tissues and were able to identify both linear and non-linear endothelial transcriptome expression changes during aging in C57BL/6 mice. We observed that on average, only 8.46% of genes have changed linearly with age across all organs. This finding aligns with previous research using multi-omics data from human samples [4, 10]. The functional enrichment of these genes identified heterogeneity in how each tissue responds to aging. The increase in immune-related GO terms across major tissues might be an indication of inflammaging. Inflammaging is considered one of the seven pillars of aging and has been widely reported during the aging process [37]. Studies have shown that lifetime or short-term dietary restriction treatments can ameliorate inflammaging in mice [38]. In our study, tissues such as the liver, kidney, heart, and gWAT displayed signs of a gradual increase in inflammation-related genes. This is not entirely surprising when we consider that these tissues are highly vascularized and would be the first responders to aging-related cues. Moreover, the liver was the only tissue, among those analyzed, that underwent drastic metabolic reprogramming as it aged. Studies have shown that downregulation of major metabolic pathways in the liver is associated with cellular senescence and low-grade inflammation, especially in hepatocytes, liver sinusoidal ECs, and Kupffer cells [39]. This trend is also evident in our analysis, as we see an upregulation of lymphocyte-mediated immunity in the ECs. The fact that we see only a small percentage of genes being affected by aging linearly gains significance in light of the recently proposed concept of gene length–dependent transcriptional decline (GLTD) [40]. GLTD represents a bottleneck in the transcriptional machinery, suggesting a negative correlation between the linear expression of genes and their gene length in the aging process. Exploring whether GLTD contributes to the limited set of linearly age-associated genes in ECs could provide an important extension to this study.

Through this study, we report for the first time the nonlinear nature of transcriptomic changes in ECs across different tissues in mice. The inclusion of five distinct age stages in our dataset enabled us to capture the wave-like, non-linear changes associated with endothelial aging in mice. Interestingly, similar non-linear dynamics in cellular composition have previously been reported using these datasets in the PanSci atlas [11]. However, in our analysis, we did not observe significant changes in the relative abundance of vascular EC subtypes across any of the tissues. Although Aristotle described aging as ‘a gradual cooling of the body’ [41], studies over the years have gradually contradicted this. For instance, it is known that during menopause, female reproductive organs undergo a significant transition [42]. Hematopoietic clonal diversity appears to decline rapidly in humans after the age of 70 years [43], and studies on the human proteome identified an abrupt increase and decrease of certain proteins in three distinct stages of human lifespan [6]. Terms such as threshold, shifts, turning points, and transition points are being increasingly used to identify the changes that occur at the molecular level during aging [44]. One of the most reliable tools to understand how these transition points are placed across the lifespan is the DESWAN analysis tool [6]. Using this, we were able to identify major gene expression transitional points in all tissues. Interestingly, none of the organs exhibited a similar transition pattern during aging, indicating that even though all vascular ECs share a broadly conserved transcriptomic profile across the body, their gene expression programs are spatially regulated in an organ-specific manner. This pattern was consistently observed across the functional roles of these genes in different organs, further supporting these findings.

With this statement, we also acknowledge the limitations of this study. 1) This data is limited by the lack of temporal resolution; i.e., the availability of only 5 age groups for analysis may mask gradual molecular changes occurring between the sampled ages. 2) Even though we were able to identify several subtypes of vascular ECs, the lack of sufficient biological replicates stopped us from analyzing these cell types further to understand the dynamics of aging in these cell types. 3) Lack of sufficient biological replicates was also the reason why we were unable to develop an organ-specific biological clock using this transcriptomic data. 4) We also acknowledge that findings at one omics level are generally not transferable to another. Studying aging is often more effective when conducted at the multi-omics level. However, it is a highly time-consuming and expensive process. 5) Finally, our conclusions are based on just the transcriptomic data; further validation of the findings at the multi-omics level is required to confirm the identified molecular transitions.

In summary, our study provides a comprehensive understanding of EC transcriptome dynamics across the lifespan, revealing organ-specific and nonlinear aging trajectories in mice. By annotating all EC subtypes and constructing the EndoAge atlas, we were able to establish a comprehensive reference resource for exploring vascular aging at cellular resolution in all 8 tissues. Our data is thus a valuable source for understanding the vascular basis of aging, and underscores the value of murine models in studying complex physiological processes like aging, which are otherwise challenging to examine in humans.

