## Supplementary Figures for "EndoAge Atlas: Non-linear dynamics of endothelial aging reveal organ-specific vascular trajectories in mice"

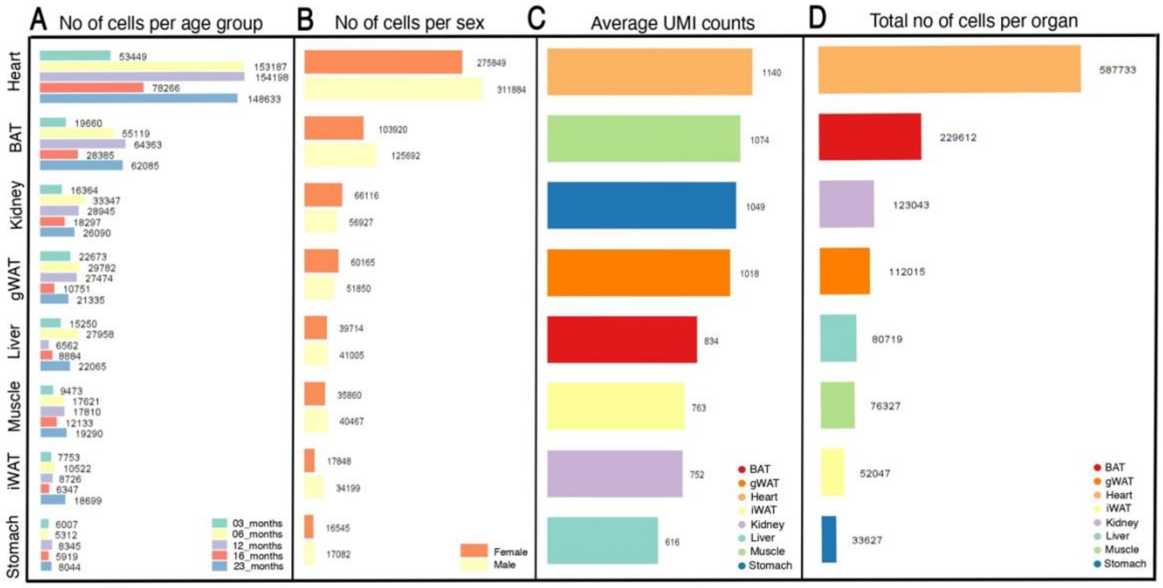


**Supplementary Fig. 1:** Barplot depicting the distribution of the number of cells across A)Ages, B) Sex across all tissues investigated. C) Barplot depicting the average UMI counts across each tissue. D) Total number of ECs detected per tissue.

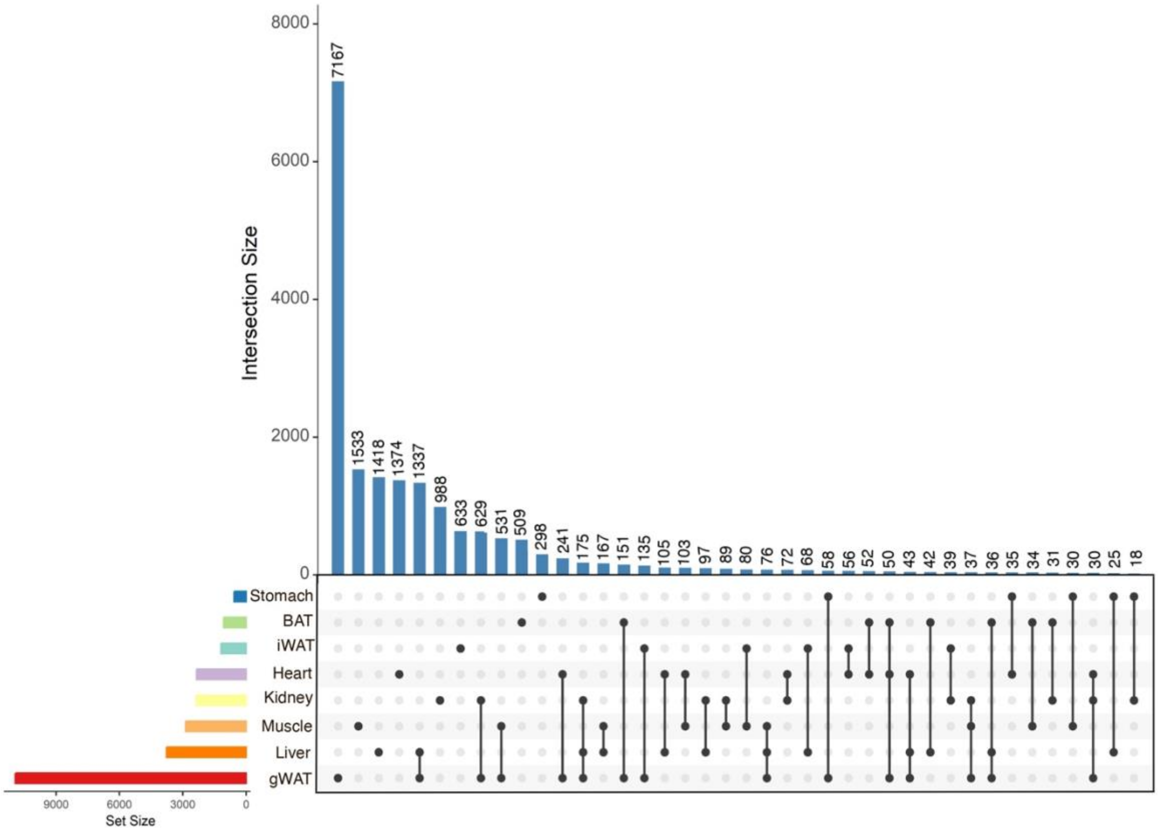


**Supplementary Fig. 2:** UpSet plot showing the number of genes exhibiting a linear correlation with age within each tissue and their overlap across tissues. Notably, no genes were found to be commonly and linearly correlated with age across all tissues.

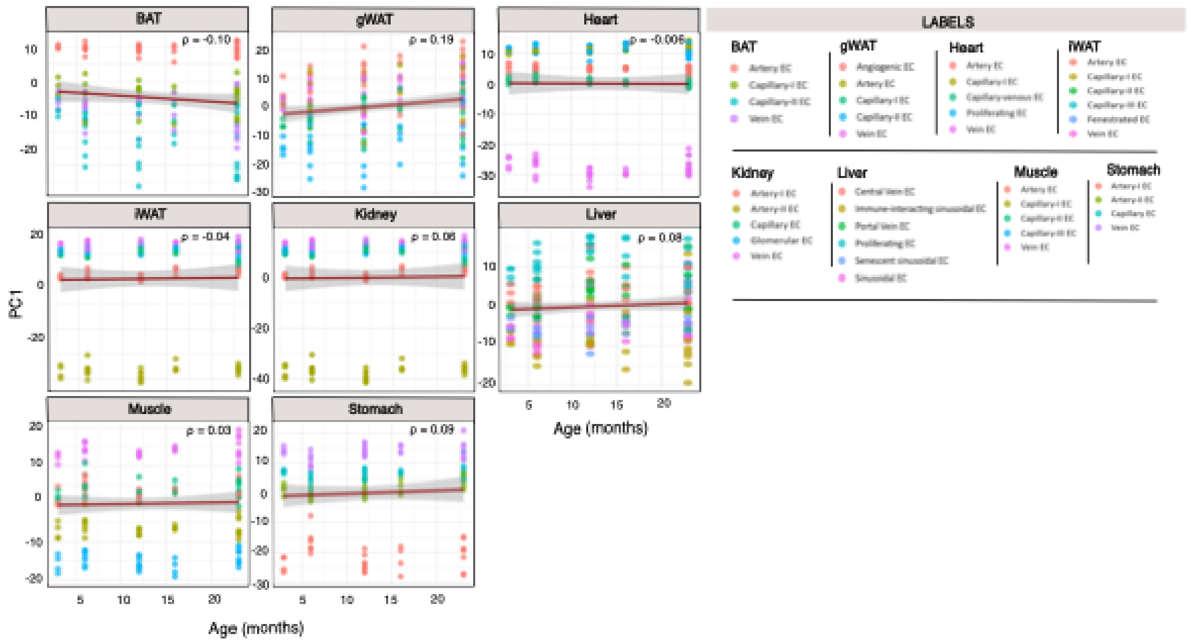


**Supplementary Fig. 3:** Scatter plot depicting the Spearman's correlation between the first principal component (PC1) and age across all tissues. Data were pseudobulked across mouse IDs and cell types, revealing cell–type–specific variation.

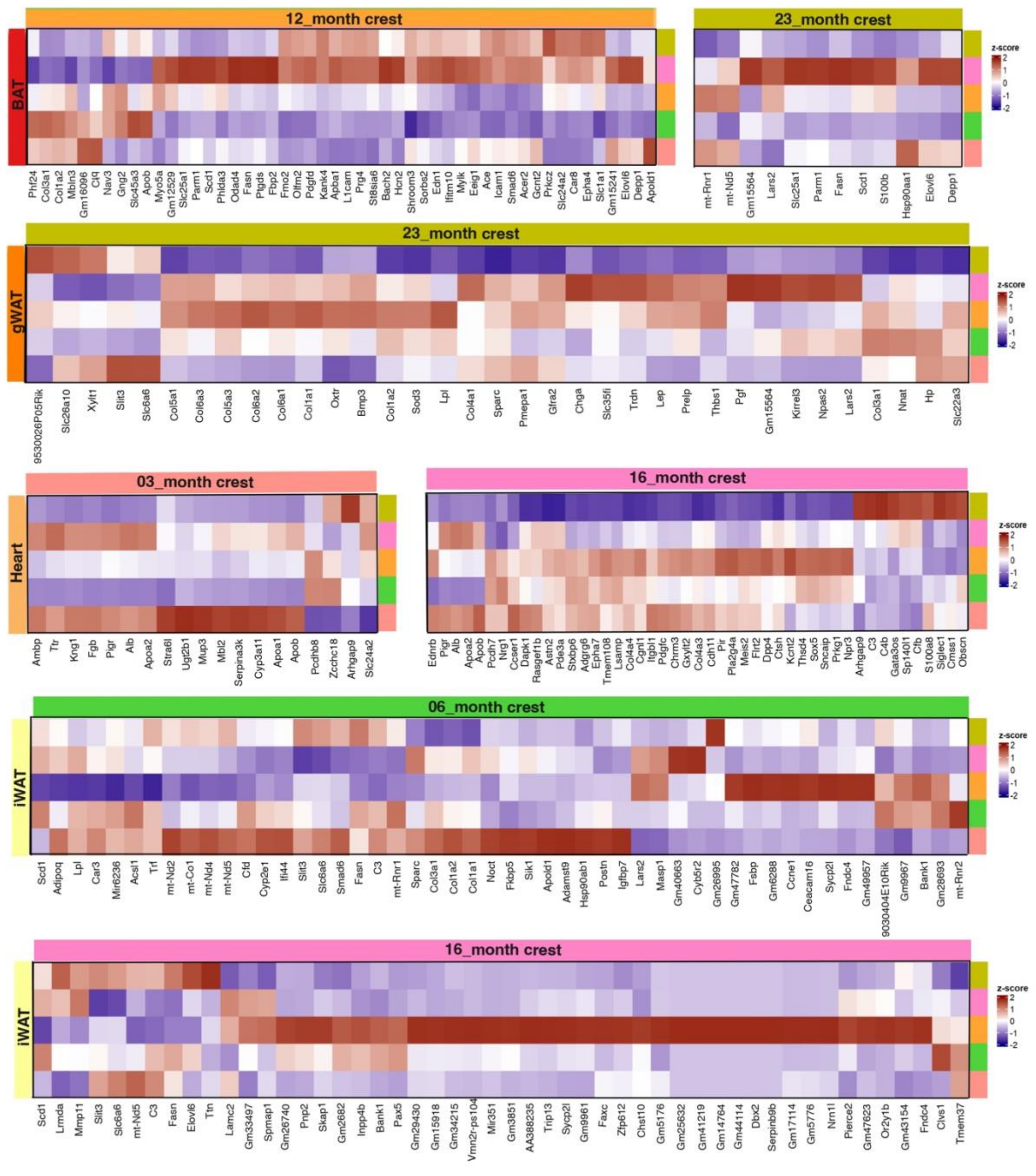


**Supplementary Fig. 4:** Heat map depicting the gene expression of the genes within the identified crests in tissues Bat, gWAT, Heart, and iWAT. Only the top 50 genes belonging to the crest in iWAT are mapped here.

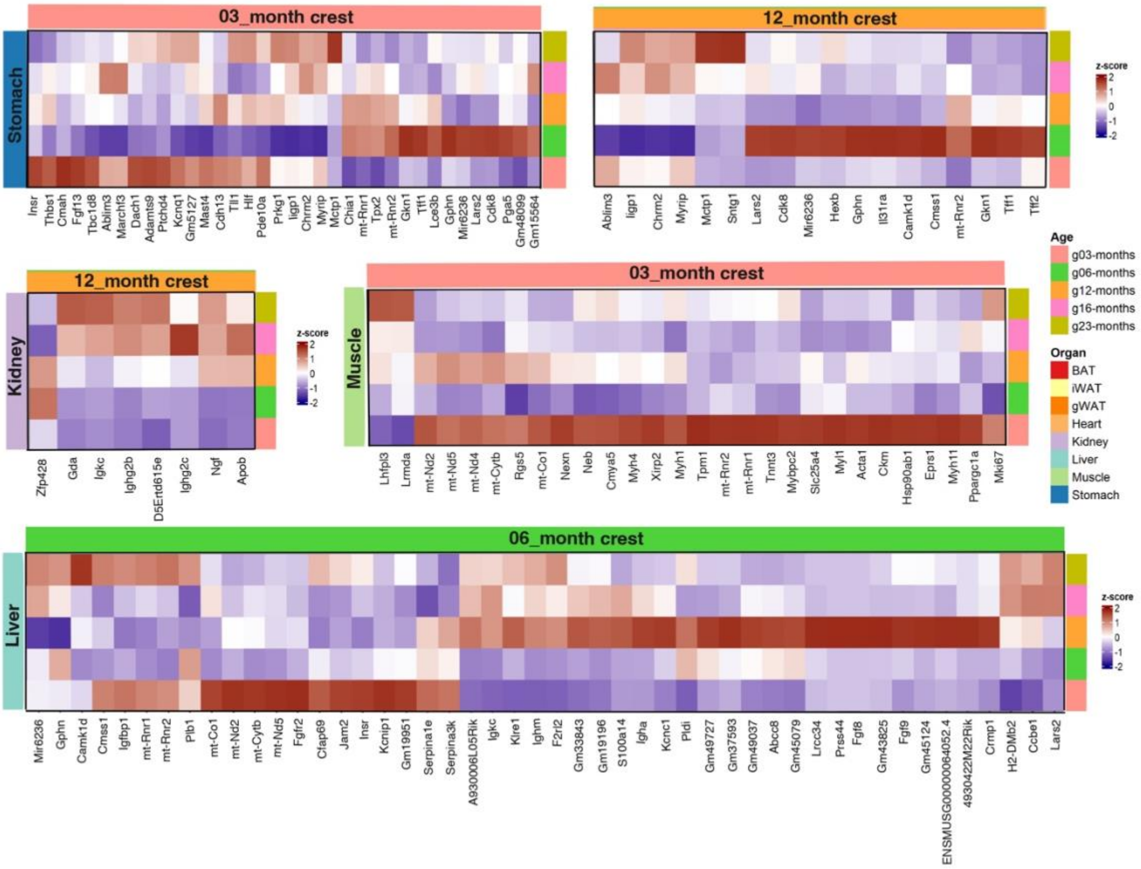


**Supplementary Fig. 5:** Heat map depicting the gene expression of the genes within the identified crests in tissues: stomach, kidney, muscle, and liver. Only the top 50 genes belonging to the crest in liver are mapped here.
